# Uncovering hidden behaviours: a GPS-Accelerometry approach to behaviour classification in storm-petrels

**DOI:** 10.64898/2025.12.23.696239

**Authors:** Darren Wilkinson, Adam Kane, Mark Jessopp, John L. Quinn

## Abstract

Biologging has transformed ecological research, yet studying the at-sea behaviour of small seabirds remains challenging due to trade-offs between device size and data resolution. Miniature GPS devices (<1 g) have been deployed on storm-petrels but their coarse temporal resolution limits behavioural classification, particularly resting. Accelerometers offer detailed behavioural data but lack spatial context, and some species are too small to simultaneously carry both devices. We deployed GPS devices on breeding European storm-petrels, including a subset recently developed with integrated accelerometers. Using accelerometer-informed rest periods, we developed a semi-supervised three-state Hidden Markov Model to classify transiting, foraging, and resting across all GPS tracks, even those without accelerometer data. Detection of resting behaviour by the semi-supervised model improved substantially compared to an unsupervised GPS-only model which only identified 17% of true resting events. Resulting time-activity budgets closely matched accelerometer-only studies, validating behaviour classifications, while also providing spatial context. Foraging peaked in the early hours of the night, whereas resting was most frequent in the afternoon. This is the first application of a combined GPS-accelerometer approach to a small storm-petrel species. Our method improves behavioural classification in small seabirds, providing new insights into the spatial and temporal dynamics of their at-sea behaviour.

## Introduction

Biologging has become an essential tool in ecological research, with applications spanning a wide range of species and ecosystems (Hussey et al., 2015; Kays et al., 2015; Sequeira et al., 2025). The deployment of devices capable of recording high-resolution spatial and temporal data has provided unprecedented insights into animal movement (Goldshtein et al., 2022), behaviour (Darby et al., 2022a), and physiology (Williams & Ponganis, 2021). Global Positioning System (GPS) devices record animal movement with high spatial accuracy (Cagnacci et al., 2010), enabling researchers to investigate species daily movement patterns (McDuie et al., 2019), migration routes (Pedersen et al., 2019), habitat use (Roeleke et al., 2016), social interactions (McIlraith et al., 2025), and predator-prey dynamics (Eriksen et al., 2011). When paired with statistical tools such as Hidden Markov Models (HMMs), GPS data can be used to infer behavioural states based on movement characteristics (Bodey et al., 2014; Farhadinia et al., 2020). However, the reliability of these inferences is limited by the temporal resolution of GPS fixes (Postlethwaite & Dennis, 2013) and the overlapping movement patterns associated with different behaviours (Bennison et al., 2019).

Accelerometers, in contrast, record acceleration along one, two, or three axes at extremely high temporal resolutions, far exceeding that of GPS devices. These data enable the identification of discrete behaviour states such as flying, running, and resting (Shepard et al., 2008) allowing the construction of time-activity budgets (Collins et al., 2016; Beltramino et al., 2019; Tatler et al., 2021; Yu et al., 2022). They also capture brief, transient behaviours (Kawabata et al., 2014), and metrics derived from dynamic acceleration have been linked to energy expenditure across a broad range of taxa (Halsey et al., 2009; Wilson et al., 2020). However, device placement position can significantly influence data quality, with optimal results achieved when placed near the centre of gravity (Wilson et al., 2020; Garde et al., 2022). Furthermore, as accelerometers lack spatial context, they cannot reveal where behaviours occur or whether they are influenced by environmental cues. For this reason, accelerometers are often deployed in combination with GPS devices (e.g. Berlincourt et al., 2015; Wilmers et al., 2017).

Historically, the size and weight of biologging devices have restricted their use to larger species due to ethical concerns regarding animal welfare (Casper, 2009), leaving significant knowledge gaps for smaller species. This is particularly true in seabird research where larger biologging devices have been shown to negatively impact some species (Bodey et al., 2018; Gillies et al., 2020; Clairbaux et al., 2025). While larger seabirds such as the wandering albatross (*Diomedea exulans*) have been studied with data loggers for decades (Jouventin & Weimerskirch, 1990), only recent advancements in device miniaturisation have made it possible to track the smallest seabirds, the storm-petrels (e.g. Hedd et al., 2018; Lago et al., 2019; Rotger et al., 2020; Bolton, 2021; Alho et al., 2022). Moreover, although the simultaneous use of GPS tags and accelerometers on seabirds is becoming more common (e.g. Berlincourt et al., 2015; Ma et al., 2022; Sutton et al., 2023), no such deployments have been achieved on species as small as storm-petrels.

The European storm-petrel (*Hydrobates pelagicus*), weighing approximately 26 g (Brooke, 2004), is the smallest seabird breeding in the North Atlantic. GPS tracking has been conducted at a limited number of colonies (e.g. Rotger et al., 2020; Bolton, 2021; De Pascalis et al., 2021), but these light-weight devices are constrained by their battery life and storage capacities. As a result, GPS data have typically been recorded at 30 min intervals (Bolton, 2021), or as infrequently as every one or two hours (Rotger et al., 2020; De Pascalis et al., 2021), and this coarse temporal resolution hinders behaviour inference. Storm-petrels have been observed resting at sea (Aguado-Giménez et al., 2016) and given that they undertake multi-day foraging trips over long distances (Rotger et al., 2020; Bolton, 2021; De Pascalis et al., 2021), such resting behaviour is likely a vital component of their energy budget. However, HMMs applied to temporally coarse GPS data have only been able to distinguish two behavioural states, interpreted as transiting and foraging (De Pascalis et al., 2021; Deakin, 2022), with resting likely misclassified as foraging (Diop et al., 2018). Accelerometers have recently provided the first insights into the flight strategy, behavioural complexity, and activity budgets of European storm-petrels in the Mediterranean (De Pascalis et al., 2025). Yet, due to weight limitations, these birds have not been tracked with GPS tags and accelerometers simultaneously, preventing the spatial context of these behaviours from being resolved. Furthermore, this lack of integration has meant that the accuracy of GPS-based HMM behavioural classifications for storm-petrels remains unvalidated using other data sources.

Recent technological advancements have enabled the development of miniaturised GPS devices with integrated accelerometers, allowing for the first multi-sensor study of the smallest seabirds. Here, we present the first application of a GPS-accelerometry approach to behaviour classification in storm-petrels, allowing examination of the spatial and temporal dynamics of at-sea behaviours that were previously inaccessible with GPS-only or accelerometer-only methods. We used accelerometer data from a small sample of foraging trips to identify rest periods, which were then incorporated to semi-supervise a Hidden Markov Model applied to the GPS data, enabling classification of three distinct behaviours (transiting, foraging, and resting) even when GPS tracks were temporally coarse and lacked accelerometer data. This approach offers new opportunities to identify where, when, and how storm-petrels encounter anthropogenic pressures at sea, and to assess how climate-driven changes in the marine environment may influence their behaviour and energy budgets. Overall, the framework presented here provides a replicable method for studying the movement and behavioural ecology of other small seabird species.

## Methods

### Data collection

A total of 35 Pathtrack nanoFix Geo-mini (<1 g) GPS devices were deployed on breeding European storm-petrels from two colonies over four breeding seasons. In 2016, eleven tags were deployed on chick-rearing birds at High Island, Co. Galway, Ireland (53.546 N, -10.258 W), while 24 devices were deployed at Illauntannig, Magharee Islands, Co. Kerry, Ireland (52.326 N, -10.022 W) across 2020, 2023, and 2024 (7 incubation, 6 chick-rearing, 11 unconfirmed breeding stage). Further details on capture methods can be found in the electronic supplementary material (ESM) Appendix A. All tags were programmed to record locations in 30 min intervals except four tags in 2024 that had a 15 min temporal resolution. These four tags also included tri-axial accelerometers that recorded data continuously at 25 Hz with a sensitivity range of ±8 g. Body mass was recorded before fitting and after removing the GPS device. In all years, the devices were attached to the four central tail feathers using Tesa tape, following standard protocols for this species. Total deployment weight represented 3.48 ± 0.27% of the bird’s body mass.

### Step 1: GPS-derived behaviours

All analyses were conducted using R version 4.5.1 (R Core Team, 2025). Hidden Markov Models (HMMs) can infer at-sea behaviour states of seabirds by analysing step lengths and turning angles derived from GPS data (e.g. Bodey et al., 2014; Bennison et al., 2018). To correct for any missed or delayed GPS fixes, tracks were linearly interpolated to consistent 30 min intervals using the *move2* package (Kranstauber et al., 2024). As a result, trips recorded at a 15 min temporal resolution were subsampled to align with the resolution of the other tracks. If gaps between GPS fixes exceeded three hours, tracks were split into sections to prevent interpolating over extended time intervals (De Pascalis et al., 2021). Tags affected by technical issues that prevented GPS data recording for most of the trip were excluded from the analysis. A 0.5 km buffer surrounding the colony was used to define the start and end of each foraging trip. A trip was considered to begin at the first recorded point outside the buffer and to end upon re-entry. All location points within the buffer, both before departure and after return, were excluded from the analysis. To determine the optimal number of states for this dataset, we constructed HMMs with one to four states using the *momentuHMM* package (McClintock & Michelot, 2018). Every increase in the number of behaviour states corresponds with an increase in log-likelihood and therefore favours complex, overfitted models. Consequently, the optimal number of states was selected based on the model that produced the largest increase in log-likelihood (Dean et al., 2013) representing a compromise between model complexity and accuracy. In all HMMs, step length and turning angle were modelled using a gamma and Von Mises distribution with a mean turning angle of zero, respectively, and starting parameters were selected using *k*-means clustering (Dean et al., 2013; Saldanha et al., 2023). The Viterbi algorithm was applied to identify the most probable sequence of behaviour states according to the best model. This process was repeated using only the subset of tracks recorded at 15 min intervals, with interpolation applied to ensure this temporal resolution was consistent across each track, to assess whether the increased resolution would enable the HMM to distinguish a greater number of behavioural states.

### Step 2: Accelerometer-derived behaviours

Several metrics that may reflect distinct behavioural states were extracted from tri-axial accelerometer data collected during three chick-rearing European storm-petrel foraging trips (two complete, one incomplete). The Z-axis (heave) is frequently used to distinguish seabird behaviours (e.g. Ropert-Coudert et al., 2006; Sakamoto et al., 2013; Patterson et al., 2019; Connors et al., 2021; Schoombie et al., 2023), and the standard deviation of the Z-axis (SD_Z_) was calculated from the raw acceleration data over a one-second moving window (Collins et al., 2015). Measures of dynamic acceleration can also be used to separate behaviours (Patterson et al., 2019) and therefore to isolate dynamic acceleration, the gravitational (static) component for each axis was estimated using a one-second moving average and then subtracted from the raw acceleration. From this, Overall Dynamic Body Acceleration (ODBA) was computed as the sum of the absolute values of dynamic acceleration across all three axes (Wilson et al., 2006). The mean and standard deviation of ODBA over 25 data points (i.e. one second) was calculated.

After visual inspection of the ODBA values for each foraging trip, it was apparent that resting behaviour (interpreted as low ODBA) was more pronounced in one of the trips compared to the others. Each metric exhibited a clear bimodal distribution for this trip (Figure S1), corresponding to two behavioural states (resting and flying). The interpeak frequency minimum, which is the value at the trough between the two modal peaks, was identified using kernel-density estimation and then verified by visual inspection. This value was used as the threshold for separating resting from flying (Collins et al., 2015). Only one of the three accelerometer datasets contained sufficient resting behaviour to produce clear bimodal distributions for any metric, enabling reliable extraction of the interpeak frequency minimum. As the other two datasets contained very limited resting and therefore did not exhibit stable bimodality, their thresholds could not be estimated independently. We therefore used the thresholds derived from the resting-rich dataset and applied it to the remaining two datasets. This approach assumes that the acceleration signatures of resting versus flying is comparable across individuals under the same device placement and sampling conditions. To confirm that this was reasonable, we visually inspected the metric distributions and verified that the transferred threshold fell at the expected low-value end of the distribution (Figure S2). All metrics were strongly correlated with one another (>0.9), so the standard deviation of ODBA (SD_ODBA_) was selected to distinguish rest from flight, as it is less sensitive to orientation changes caused by tail movements that can bias individual axis measurements (Halsey et al., 2011). The three accelerometry tracks were segmented into periods of “resting” and “flying”. Resting was defined as any interval of at least 20 seconds during which ≥70% of SD_ODBA_ values remained below the interpeak frequency minimum (Figure 1). A 70% threshold was used to accommodate occasional tail movements that may occur while resting (see ESM Appendix B for sensitivity analysis). In contrast to back-mounted accelerometers (De Pascalis et al., 2025), the tail-mounted devices used in this study did not allow for reliable differentiation between dynamic behaviours such as transiting and foraging.

**Figure 1.**
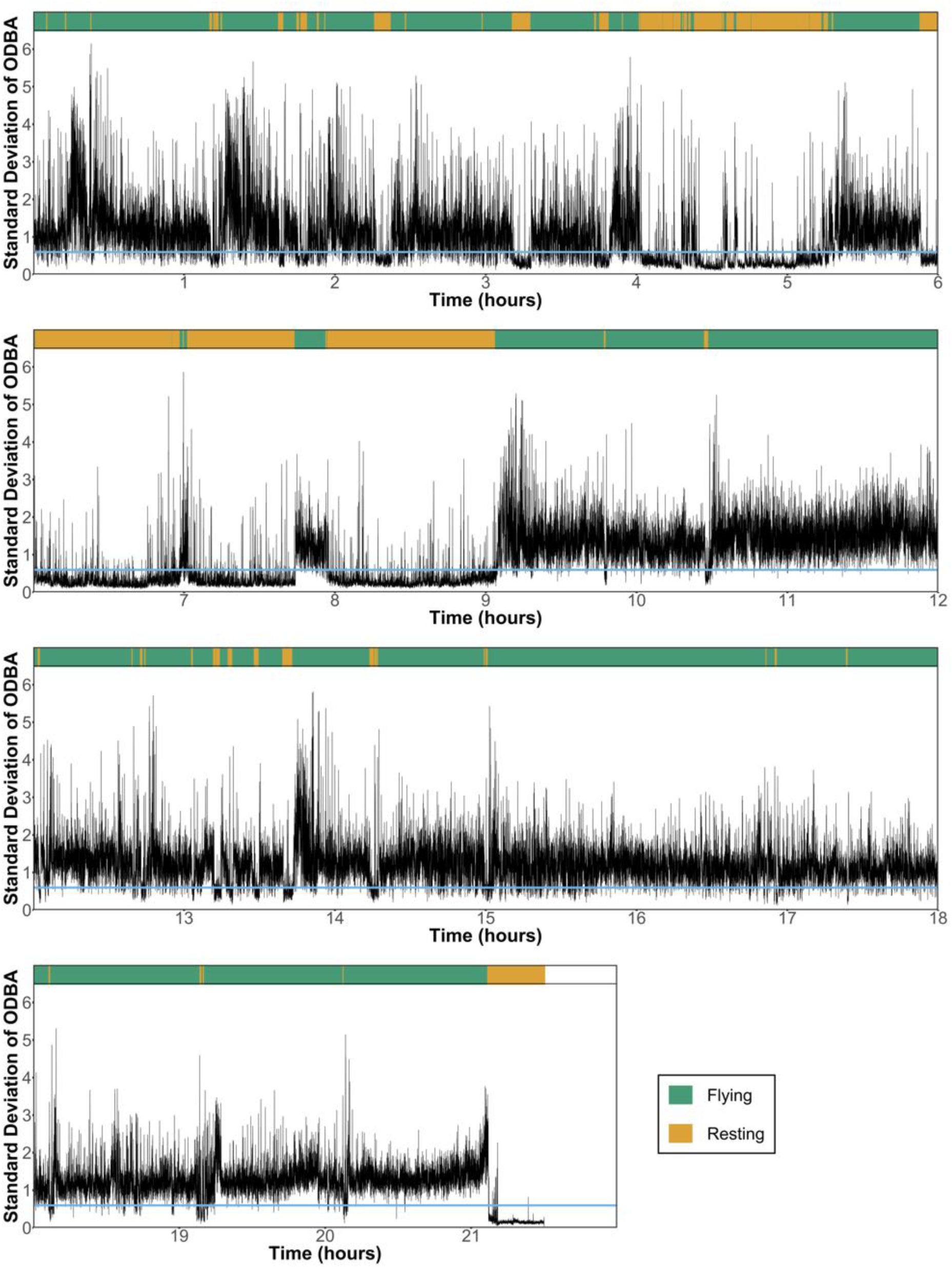
Standard deviation of Overall Dynamic Body Acceleration (ODBA) calculated over a one second moving window for a European storm-petrel foraging trip. The top bar shows periods identified as “flying” (green) and “resting” (orange). The blue horizontal line marks SD_ODBA_ = 0.597, the interpeak frequency minimum (Figure S1) used to separate the behaviours.

### Step 3: Informing Hidden Markov Models with accelerometer data

The accelerometer-informed behaviours for these three tracks were used to refine the behaviour states predicted for them by the base HMM (two-state HMM; see results) created using the methods outlined in Step 1. As the GPS data were recorded at 30 min intervals, each GPS point was classified as “resting” if the accelerometer indicated resting behaviour ≥50% of the preceding 30 min period. If resting behaviour was detected for <50% of that interval, the GPS point retained its original classification from the base HMM as either “foraging” or “transiting”. These labelled behaviours were then used to produce a semi-supervised three-state HMM by incorporating them using the *knownStates* argument within the ‘fitHMM’ function from the *momentuHMM* package (McClintock & Michelot, 2018). In practice, this means that the model was given the true behavioural states (transiting, foraging, and resting) for the three tracks that included accelerometer data allowing the model to learn the characteristic movement signatures (step lengths and turning angles) associated with each behaviour and estimate more accurate state transition probabilities. The semi-supervised HMM used these learned movement patterns to classify behaviours in the remaining GPS-only tracks. The starting parameters for the HMM were selected by evaluating the log-likelihood values from 50 candidate models, each run with randomly selected priors for step length and turning angle concentration from a range of reasonable values (De Pascalis et al., 2021). A gamma distribution for step length and a Von Mises distribution with a mean of zero for turning angle were used for all candidate models. The sequence of behaviour states was defined using the Viterbi algorithm. Behavioural classifications were then compared between the informed three-state HMM and the three-state HMM based solely on GPS data produced during Step 1 to check for improved accuracy. Additionally, to compare our results to those obtained using back-mounted accelerometers (De Pascalis et al., 2025), we calculated the proportion of time spent in each behaviour state during the complete foraging trips and examined the temporal distribution of behaviours defined by the three-state HMM.

## Results

### Tag retrieval

Over the four breeding seasons, 21 GPS tags were recovered but retrieval rates varied between years (ESM Appendix A). No significant difference in body mass was recorded after tag retrieval (ESM Appendix C). Four devices, including one with an integrated accelerometer, experienced technical faults that led to a failure to record GPS locations for most of the trips, and were therefore removed from the analyses. The remaining 17 devices resulted in 11 complete and 8 incomplete trips. Two tags deployed at the Magharee Islands colony recorded two complete trips each as the storm-petrels were not captured on their first return to the colony, while the other tags recorded one trip each.

### Behaviour state classification

A two-state model, which inferred behaviours as “foraging” and “transiting”, was selected as the base HMM for further analysis, as the largest increase in log-likelihood was identified between the one- and two-state models (Table S1). The same result was obtained using just the tracks recorded with a 15 min temporal resolution.

The accelerometers programmed to record within a range of ±8 g rarely reached this limit in any of the three axes (<0.001%). These data detected resting periods in all three tracks (two complete and one incomplete), though the extent of resting varied between individuals. According to the accelerometer data, one European storm-petrel that completed a one-day foraging trip rested for 25.2% of the time, while the other complete trip and incomplete trip revealed the birds rested for 2.0% and 2.8% of the time, respectively. The longest continuous resting period recorded was 66.7 mins, but the average duration was just 3.3 mins. The histogram of SD_ODBA_ for the foraging trip containing a substantial proportion of resting behaviour showed a bimodal distribution with an interpeak frequency minimum of 0.597 (Figure S1).

Using the periods of resting identified by the accelerometers to update the behaviour states defined by the base HMM for these three tracks resulted in the “foraging” state being split into 96 foraging and 12 resting points. All 39 points classified as “transiting” by the two-state HMM retained this status as the accelerometers never recorded close to the 50% threshold used to indicate resting behaviour for these points. Using these as “known states” to supervise the HMM allowed for the identification of transiting (characterised by large step lengths and low turning angles), foraging (characterised by intermediate to short step lengths and high turning angles), and resting (characterised by very short step lengths and moderate turning angles) in all GPS tracks (Figure 2; Figure 3; see ESM Appendix D for more details). Out of the 11 complete trips performed by breeding European storm-petrels, the three-state HMM informed by accelerometer data identified resting behaviour in 10 trips. The single trip in which no resting was detected was relatively short (19 hours), making it plausible that resting behaviour did not occur. There was great individual variation in the proportion of time allocated to each behaviour during complete trips (Figure S3) but on average 85.5% of trips was spent flying split between transiting (41.3%) and foraging (44.2%). Resting only accounted for an average of 14.5% of the time spent at sea but with great individual variation (0-31%, n = 11). Resting behaviour was more frequent during the day, peaking between 13:00 and 14:00 (Figure 4b). Foraging activity peaked in the early hours of the night (Figure 4a), whereas transiting was most common in the hours following sunrise (Figure 4c).

**Figure 2.**
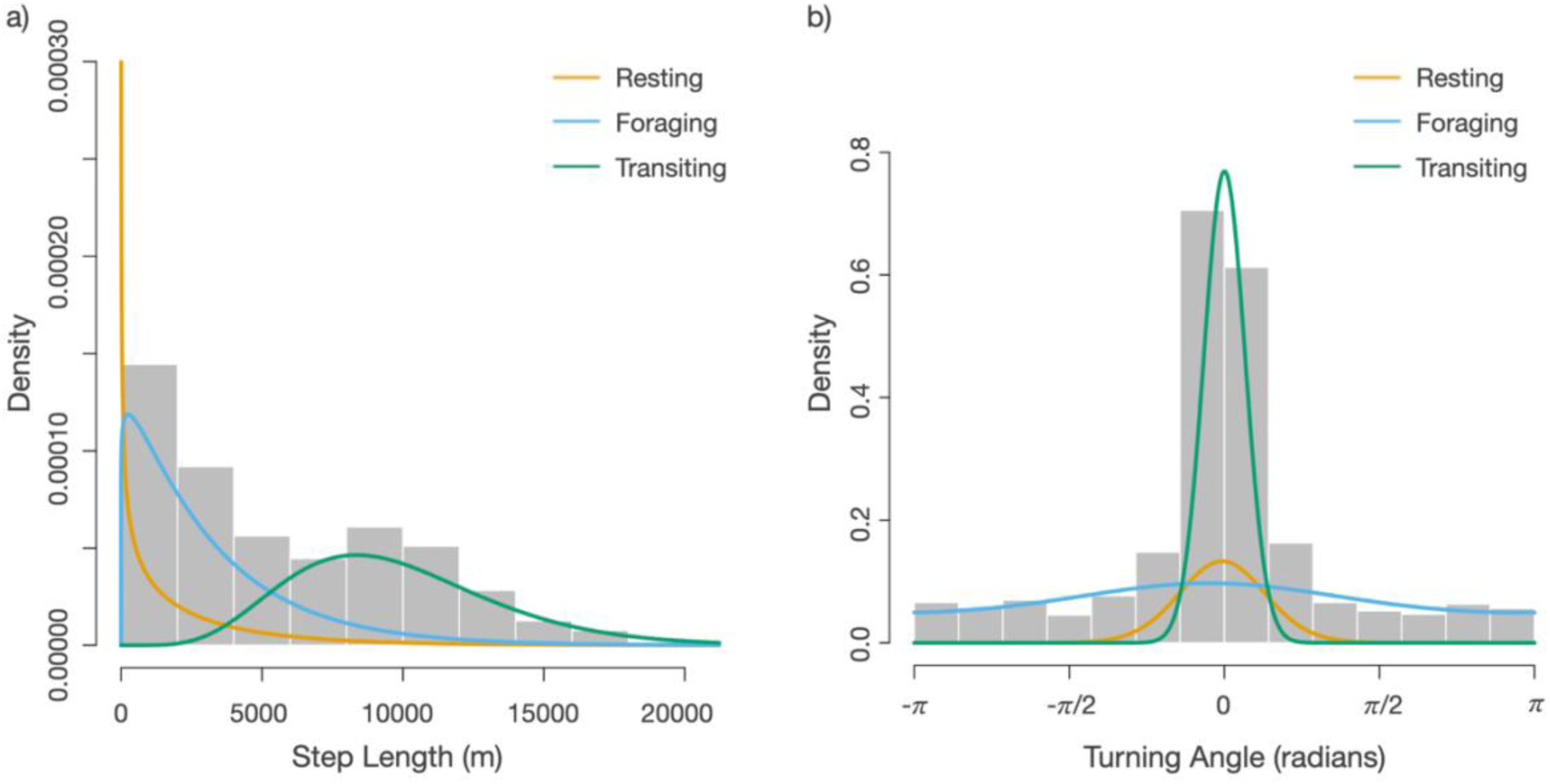
Density distributions of (a) step length and (b) turning angle of GPS data from European storm-petrels. Solid lines show the estimated state-dependent probability of three behaviour states (resting, foraging, and transiting) from the HMM, informed by the accelerometer data. See Figure S3 for individual foraging tracks segmented into each behaviour.

**Figure 3.**
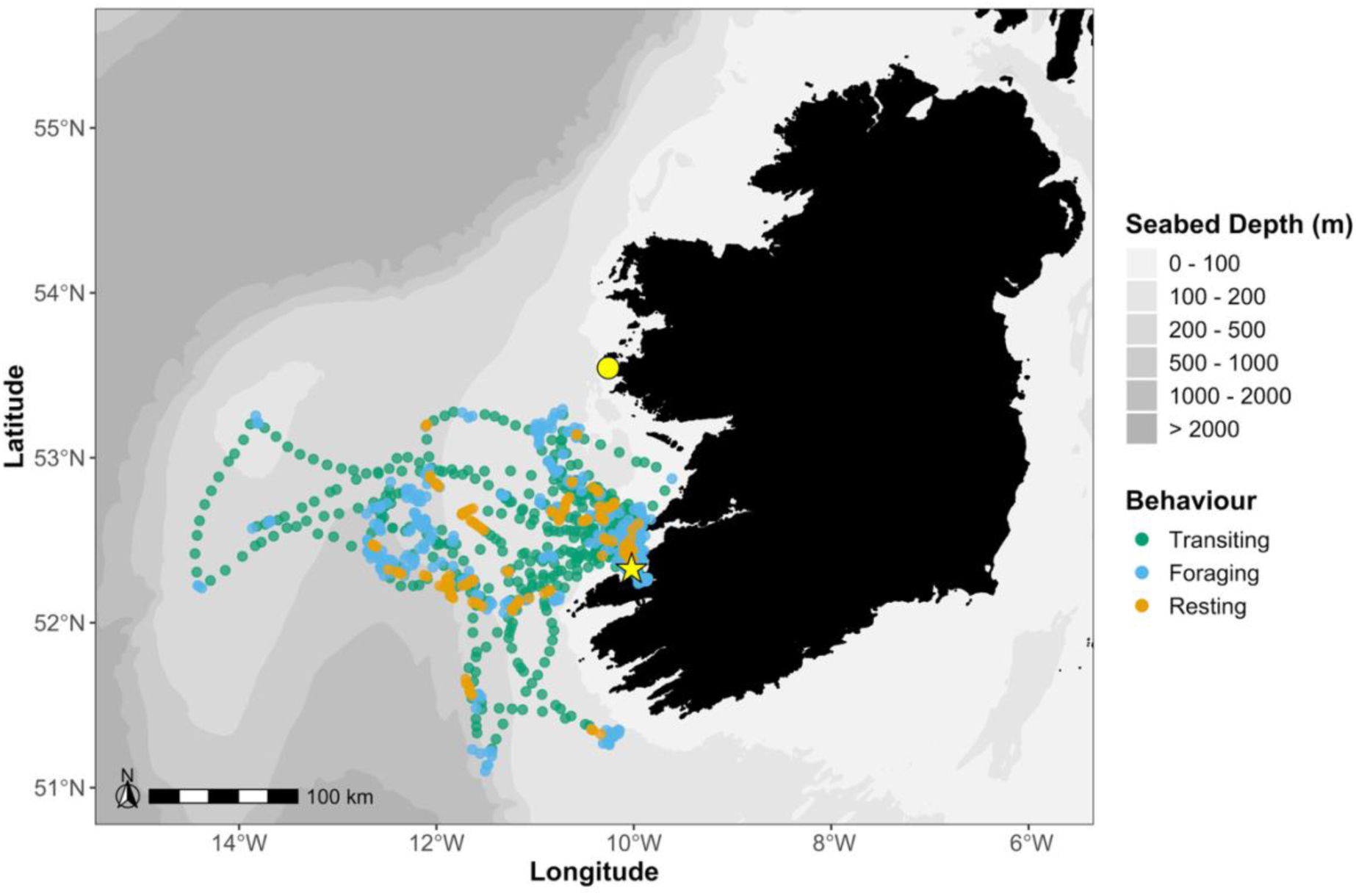
Distribution of behaviours (blue = foraging, orange = resting, green = transiting) in the 11 complete trips identified by the three-state HMM informed by accelerometer data. Yellow circle = High Island; Yellow star = Illauntannig, Magharee Islands.

**Figure 4.**
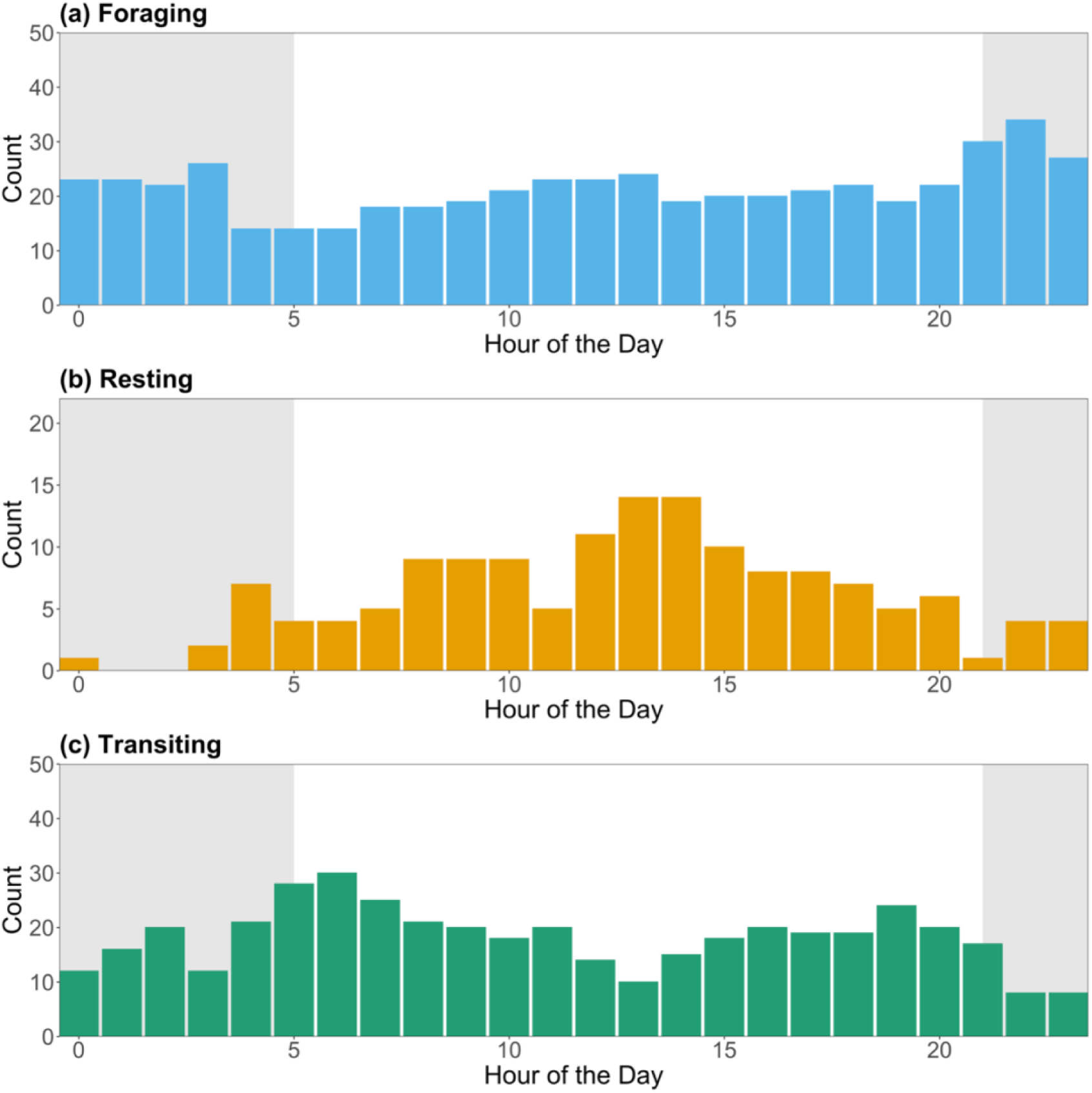
Daily activity patterns of all European storm-petrels with complete GPS-tracked foraging trips (n = 11). The grey shaded areas indicate night-time.

By comparing the behavioural classifications produced by the semi-supervised HMM, which incorporated accelerometer-informed behaviours, with those from the three-state GPS-only HMM developed in Step 1, we found strong agreement with 85% of all points matching behaviour classification. This suggests that imposing a three-state structure on the GPS-only dataset, despite the earlier selection of a two-state model as optimal, could provide reasonably accurate behavioural classification for storm-petrels. However, there were substantial discrepancies for resting. Of the 659 points labelled as transiting by the accelerometer-informed model, 93% were also identified as transiting by the GPS-only HMM (Table S2). Agreement was even higher for foraging, with the GPS-only model matching 99% of the 727 foraging points identified by the informed model (Table S2). In contrast, the GPS-only model struggled to detect resting behaviour, classifying just 4% of points across all complete and incomplete trips as resting compared to 14% of points by the informed model. Of the 219 resting points classified by the accelerometer-informed HMM, only 17% were also recognised as resting by the GPS-only model, with the majority of remaining points misclassified as foraging (Table S2).

## Discussion

This is the first study to classify the at-sea behaviour of any storm-petrel species using an integrated GPS-accelerometry approach. By combining both data sources we were able to overcome key limitations of GPS-only models and developed a semi-supervised HMM capable of distinguishing transiting, foraging, and resting behaviours. This methodological advance provides new opportunities not only to improve behavioural classification in small seabird species but also to enhance understanding of how these species allocate time and energy at sea.

The selection of a two-state HMM, separating transiting and foraging behaviours, when using only GPS data aligns with Deakin (2022) and De Pascalis et al. (2021) but it underestimates the behavioural complexity of European storm-petrels which are also known to rest at sea (Aguado-Giménez et al., 2016; De Pascalis et al., 2025). Distinguishing resting from foraging behaviour using GPS data alone is challenging (Diop et al., 2018; Bennison et al., 2019) as both behaviours exhibit short step lengths, increasing the likelihood of misclassification. Many seabird tracking studies employing HMMs benefit from high-frequency GPS data (2-10 mins) allowing the identification of three or more behavioural states (e.g. Grecian et al., 2018; Trevail et al., 2019; Darby et al., 2021). In contrast, even with 15 min GPS intervals, a two-state model remained the best fit for European storm-petrels indicating that finer-scale data are necessary to reliably separate resting from foraging behaviour. However, current GPS technology suitable for this small species lacks the battery capacity to support such high-resolution tracking over their multi-day foraging trips.

Continuous accelerometer data allow for more accurate behaviour classification and time-activity budget estimation than GPS data alone (Yu et al., 2022). Accordingly, the most comprehensive insights into the at-sea behaviour of European storm-petrels to date have come from accelerometry. De Pascalis et al. (2025) used back-mounted accelerometers to identify three behaviours across seven individuals: transiting (42%), foraging (36%), and resting (22%). Placing devices on the back is considered optimal for estimating energy expenditure (Wilson et al., 2020) and therefore for the most accurate behaviour classification when using energetic metrics (Patterson et al., 2019), as this location is close to the bird’s centre of gravity. However, when accelerometers are deployed without GPS devices, behaviours lack spatial context. In contrast, our study employed tail-mounted GPS tags with integrated accelerometers, providing simultaneous spatial and acceleration data. These devices were tail-mounted, rather than back-mounted, to ensure secure attachment and prevent the long external antenna from being obstructed in the bird’s plumage or causing discomfort. Although tail movements may introduce uncertainty in behaviour classification, clear bimodal distributions in SD_Z_, mean ODBA, and SD_ODBA_ (Figure S1) enabled robust separation of high-activity (flying) from low-activity (resting) behaviours. Resting periods were typically brief, lasting only a few minutes, and varied between individuals likely explaining why HMMs based solely on GPS data have struggled to detect resting. By incorporating accelerometer-informed rest periods into the HMM, we developed a semi-supervised three-state model capable of distinguishing transiting, foraging, and resting across all trips.

However, several limitations of our study should be acknowledged. First, our analyses were based on a relatively small sample of GPS tracks (11 complete, 8 incomplete) and only three accelerometers, only one of which provided sufficient resting periods to reliably identify the interpeak frequency minimum used to separate high-activity flying from low-activity resting. Although studies with limited sample sizes can yield valuable insights, their generality is constrained. Seabird tracking studies frequently report colony-specific foraging distributions (e.g. Wakefield et al., 2013; Bolton et al., 2019; Kane et al., 2020) and substantial individual variation in foraging behaviour (e.g. Kato et al., 2000; Patrick et al., 2015; Phillips et al., 2017; Krüger et al., 2019), which may not have been fully captured here. Device placement may also have influenced results. Accelerometers ideally should be positioned near the centre of gravity (Wilson et al., 2020), and tail placement made it impossible to distinguish foraging from transiting. By identifying 20 min periods during which ≥70% of SD_ODBA_ values were below the interpeak frequency threshold helped filter tail-movement artefacts during resting (see ESM Appendix B), but some ambiguity likely remains. Finally, there was evidence that autocorrelation was not fully accounted for (ESM Appendix D). While this may be mitigated through the inclusion of covariates (e.g. Klappstein et al., 2023), alternative GPS and accelerometer processing methods (e.g. Tintle et al., 2025), or different HMM structures (e.g. Stoyle et al., 2025), residual autocorrelation is expected to have only a minimal influence on behavioural state decoding.

Resting was most likely to occur during daylight hours, peaking between 13:00 and 14:00 (Figure 4b). This is consistent with De Pascalis et al.’s (2025) accelerometer-only study, though the peak occurred slightly later in ours. This discrepancy may result from factors such as regional differences in daylight hours, variation in the temporal availability of prey, or the hotter temperatures in the Mediterranean which may impose thermoregulatory stress around midday, preventing continuous flight during this time (Guillemette et al., 2016). Rotger et al. (2020) estimated that European storm-petrels spend an average of 35% of their foraging trips resting based on flight speeds calculated from GPS fixes taken at 1-2 hour intervals, while De Pascalis et al. (2025) determined resting to account for 22% of foraging trips. Our results suggest that resting time is slightly lower, averaging around 15%, though it varied considerably between individual trips (range: 0-31%, n = 11). The variation in resting time across trips, as seen in other seabirds (e.g. Collins et al., 2016), suggests that European storm-petrels may manage their energy budgets by allocating more time to resting on certain trips while limiting or even foregoing rest on others, perhaps for reasons similar to the differential needs of parents and offspring driving the dual foraging strategies seen in other seabirds (e.g. Wischnewski et al., 2019). Among the two birds for which consecutive foraging trips were recorded, one rested for only 3% of a 70 hour trip (∼2 hours) but increased resting to 10.5% on the subsequent 43 hour trip (∼4.5 hours). The other rested for 23% of a 46 hour trip (∼10.5 hours) but did not rest at all during the 19 hour trip that followed. This finding has important implications for understanding storm-petrel energetics. Although resting is often viewed as an energy-saving behaviour, it incurs its own energetic costs (Tremblay et al., 2022). Accurate quantification of individual resting behaviour is therefore essential for reliable estimates of energy expenditure.

We observed a peak in transiting behaviour around sunrise (Figure 4c), likely reflecting the time when many storm-petrels return to sea after leaving their colony which they only visit at night to reduce predation risk. This peak coincided with a decline in foraging activity, which then remained relatively consistent throughout the day (Figure 4a). A slight increase in foraging at night may reflect elevated prey availability due to the diel vertical migration of many mesopelagic species, which move closer to the surface after dark (Hays, 2003). Nocturnal foraging is a known strategy adopted by petrels and other Procellariiformes (Thomas et al., 2006; Phalan et al., 2007; Dias et al., 2012; Rubolini et al., 2015; Dias et al., 2016; Ramos et al., 2016) and is likely facilitated by daytime resting, potentially enabling individuals to conserve some energy to maximise foraging efficiency at night.

The similarity in transiting time between De Pascalis et al. (2025) and our study (42% and 41%, respectively) supports the reliability of this behavioural classification, but we found a slightly higher proportion of time spent foraging (44%) compared to 36% detected by De Pascalis et al. (2025). While this may reflect misclassification between resting and foraging by the HMM (Diop et al., 2018), ecological differences between the Atlantic and Mediterranean populations could also contribute. These include variation in storm-petrel diet (D’Elbée & Hémery, 1998; Albores-Barajas et al., 2011), weather conditions, and breeding stage as only incubating birds were tagged in the Mediterranean, while all breeding stages were represented in this study. Seabirds typically exhibit increased energy expenditure during chick-rearing compared to incubation (e.g. Collins et al., 2016), as they must meet their own energetic needs while also provisioning their chicks, likely resulting in an increase in foraging activity.

The enhanced understanding of the spatial distribution and temporal trends of behaviours provided by the methods presented in our study can play an important role in the conservation of small seabirds. These insights can help identify where and when individuals are most vulnerable to anthropogenic threats such as marine pollution and fisheries bycatch (De Pascalis et al., 2022; Clark et al., 2023; Ramírez et al., 2024), as well as how climate-driven changes in prey distribution and availability (Daunt & Mitchell, 2013) may affect European storm-petrels. Shifts in prey depth or abundance have the potential to disrupt nocturnal foraging efficiency and alter the temporal structure of behaviours, including the timing and duration of resting. Understanding these diel behavioural patterns therefore provides essential baseline information for assessing how storm-petrels may respond to ongoing ocean warming and ecosystem change.

Despite methodological and ecological differences between our study and De Pascalis et al. (2025), both studies revealed similar temporal trends and comparable time-activity budgets. This consistency supports the validity of our method and highlights the value of combining accelerometry with GPS data in small seabirds. While the results demonstrate that HMMs based on coarse GPS data can effectively distinguish transiting from shorter step length behaviours in storm-petrels, the identification of resting behaviour requires the integration of accelerometer data. While tail-mounted accelerometers may not be ideal for detailed energetic analyses, our study nonetheless provides a more detailed understanding of the at-sea behaviour of European storm-petrels compared to previous GPS studies and offers a practical framework for improving behavioural state classification in existing and future tracking datasets for this and other small seabird species.

## Conclusions

Multi-sensor studies provide detailed insights into the behaviour and ecology of seabirds, enabling the comprehensive examination of complex activities (e.g. Weimerskirch et al., 2005; Darby et al., 2022b). Our findings underscore the advantage of multi-sensor approaches to better understand the movement ecology of small pelagic species such as the European storm-petrel. Specifically, the integration of GPS and accelerometer data in our study has enabled the classification of resting behaviour, a behaviour previously undetectable with GPS tracking alone in this species. As biologging technology continues to advance, further refinement of these methods will enhance our ability to link behaviour with environmental drivers, ultimately informing more effective conservation strategies for this and other small seabird species.

## Acknowledgements

We would like to thank Manon Clairbaux, Jamie Darby, Astrid Dedieu, Micheál Fitzgerald, Luke Harman, Declan Manley and Emma Murphy for their assistance with fieldwork on Illauntannig. On High Island, we are grateful to Ash Bennison, Jodie Crane, Emma Critchley, Ronan O’Sullivan and Saskia Wischnewski for their support. Special thanks to Bob Goodwin and Feichín Mulkerrin for granting access to Illauntannig and High Island, respectively, and to Jamie Knox and Fechín Mulkerrin for facilitating transport to and from the islands. We also thank Dr Mark Bolton for his guidance on the GPS tag deployment methodology.

## Ethics

All fieldwork was approved by University College Cork’s animal ethics committee and licensed by the British Trust for Ornithology and the National Parks and Wildlife Service.

## Data Availability

GPS data used for this publication are available from the Seabird Tracking Database (https://data.seabirdtracking.org/) study numbers 1576 and 2449. All code is freely available at Zenodo (doi:10.5281/zenodo.17751693)

