## Supplementary Material for "Uncovering hidden behaviours: a GPS-Accelerometry approach to behaviour classification in storm-petrels"

---

### Supplementary Material

#### *Appendix A: Tag Recovery Rates*

Over the course of the study, 21 out of 35 GPS tags (60%) were successfully retrieved. However, recovery rates varied between years.

- 2016: 7/11 (63%)
- 2020: 5/6 (83.3%)
- 2023: 1/5 (20%)
- 2024: 8/13 (61.5%)

Despite using the same capture and recapture methods in both 2020 and 2023 (nets draped over the dry-stone walls enclosing the nests), recovery success differed. Several factors may have contributed to the low recovery rate in 2023:

1. Windy conditions on several retrieval nights in 2023 may have made the nets more visible, deterring birds from returning to their burrows.
2. By 2023, the nets had been used extensively for catching storm-petrels and as a result had a strong scent. Given this species has good olfactory senses, the storm-petrel scent outside their burrows introduced by the nets may have discouraged some birds from re-entering their nests.
3. Properly positioning nets over uneven stone walls is difficult. Loose areas can create pockets that trap birds, while taut sections may prevent entanglement, reducing capture efficiency.

In contrast, the nests used at High Island in 2016 and the Magharee Islands in 2024 were accessible, allowing birds to be captured by hand. This is the approach we recommend for future deployments where possible.

***Appendix B: Sensitivity Analysis***

Tail-mounted accelerometers are sensitive to occasional tail movements even during resting. These movements can introduce brief spikes in $SD_{ODBA}$ that fragment resting periods if smoothing is insufficient. To identify an appropriate combination of window size (W) and threshold (T) for segmenting the accelerometer data into periods of resting and flying behaviour, we conducted a multi-step sensitivity analysis. Resting was defined as any interval of at least W seconds during which ≥T of $SD_{ODBA}$ values remained below the inter-peak frequency minimum of the $SD_{ODBA}$ distribution. The goal was to select parameters that (i) produced biologically plausible patterns, (ii) are robust to small perturbations in parameter values, and (iii) minimise fragmentation of resting bouts caused by occasional tail movements.

We computed the total time resting, the total number of resting bouts, and the median resting bout duration for the accelerometer data of the European storm-petrel trip consisting of sufficient resting behaviour across a range of window sizes (5-60 s) and thresholds (0.5-0.9). Figure B1 shows heatmaps of these metrics revealing a clear structure:

- Short windows (5-10 s) and low thresholds (0.5-0.6) produce very high total resting duration, likely overestimating rest. As the window increases, total resting duration declines gradually, stabilising around 20-40s. Very high thresholds (>0.85) cause a sharp reduction in resting duration, indicating the classification becomes overly conservative.
- Very short windows detect many resting bouts (high fragmentation). Increasing the window size rapidly reduces the number of bouts, stabilising at 20-30 s, and remaining low thereafter.
- Short windows produce short median bout durations (expected due to fragmentation). Increasing window length increases median bout duration.

The combination of **W = 20 s** and **T = 0.7** fell within a stable region and avoided extremes produced by either under- or over-smoothing.

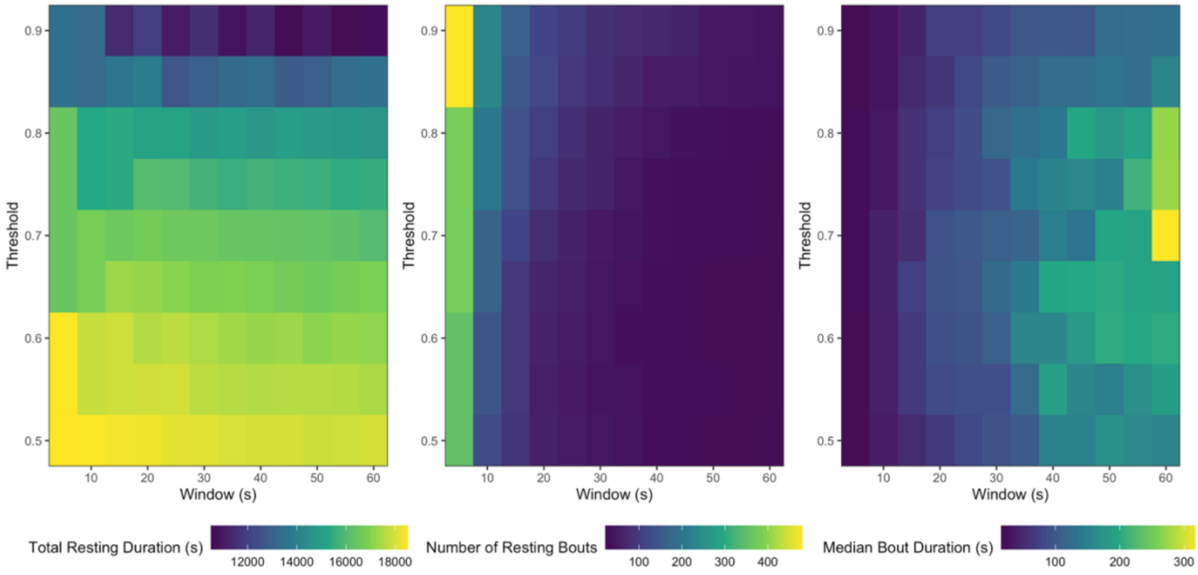


**Figure B1.** Heatmaps of resting-classification sensitivity across window sizes and thresholds

To assess robustness, we compared the resting classifications produced by the selected parameters (W = 20, T = 0.7) with those from neighbouring parameter combinations using the Jaccard similarity index. We evaluated all combinations with windows ± 5 s and thresholds ± 0.05. All neighbouring parameter combinations showed very high similarity (Jaccard = 0.91-0.97; Table B1), indicating that the classification produced by W = 20, T = 0.7 is highly robust to small changes in parameter values.

**Table B1.** Jaccard similarity index between the selected resting-classification parameters (W = 20, T = 0.7; highlighted in bold) and neighbouring parameter combinations.

| W | T | Jaccard |
|---|---|---|
| 15 | 0.65 | 0.96 |
| 15 | 0.70 | 0.96 |
| 15 | 0.75 | 0.91 |
| 20 | 0.65 | 0.97 |
| **20** | **0.70** | **-** |
| 20 | 0.75 | 0.96 |
| 25 | 0.65 | 0.97 |
| 25 | 0.70 | 0.97 |
| 25 | 0.75 | 0.96 |

#### ***Appendix C: Impacts of tagging***

We evaluated the effects of GPS tagging on breeding European storm-petrels by assessing differences in body mass pre- and post-tag deployment, and nest abandonment rates between tagged and control nests.

*Body Mass*

All European storm-petrels were weighed both before tag deployment and immediately following tag removal. In 2016, 6 out of 11 storm-petrels gained weight during deployment, while only two lost weight. We analysed pre- and post-deployment body mass using a paired t-test, incorporating data from 2020, 2023, and 2024. No significant change in body mass was found (pre-deployment mean ± SD: 27.81 ± 2.18 g; post-deployment: 27.87 ± 2.02 g; $t_{11}$ = 0.07, p = 0.94). However, body mass changes in this species are difficult to interpret due to variability in the amount of food carried for the chick.

*Nest Abandonment*

In 2024, nest abandonment was monitored between 11 experimental burrows (with GPS-tagged adults) and 12 control burrows (no handling). Of the experimental burrows, 7 (64%) remained active, 3 (27%) were abandoned following deployment (all incubating birds), and 1 (9%) had unknown status as we had to leave the colony before tag retrieval was possible. Among control burrows, 7 (58%) remained active and 5 (42%) were abandoned (all during incubation) by our final visit to the colony. While the abandonment rate was similar between experimental and control burrows, interpretation is limited by the fact that monitoring the control nests still involved some, albeit minimal, disturbance that could not be measured.

#### *Appendix D: Pseudo-Residuals*

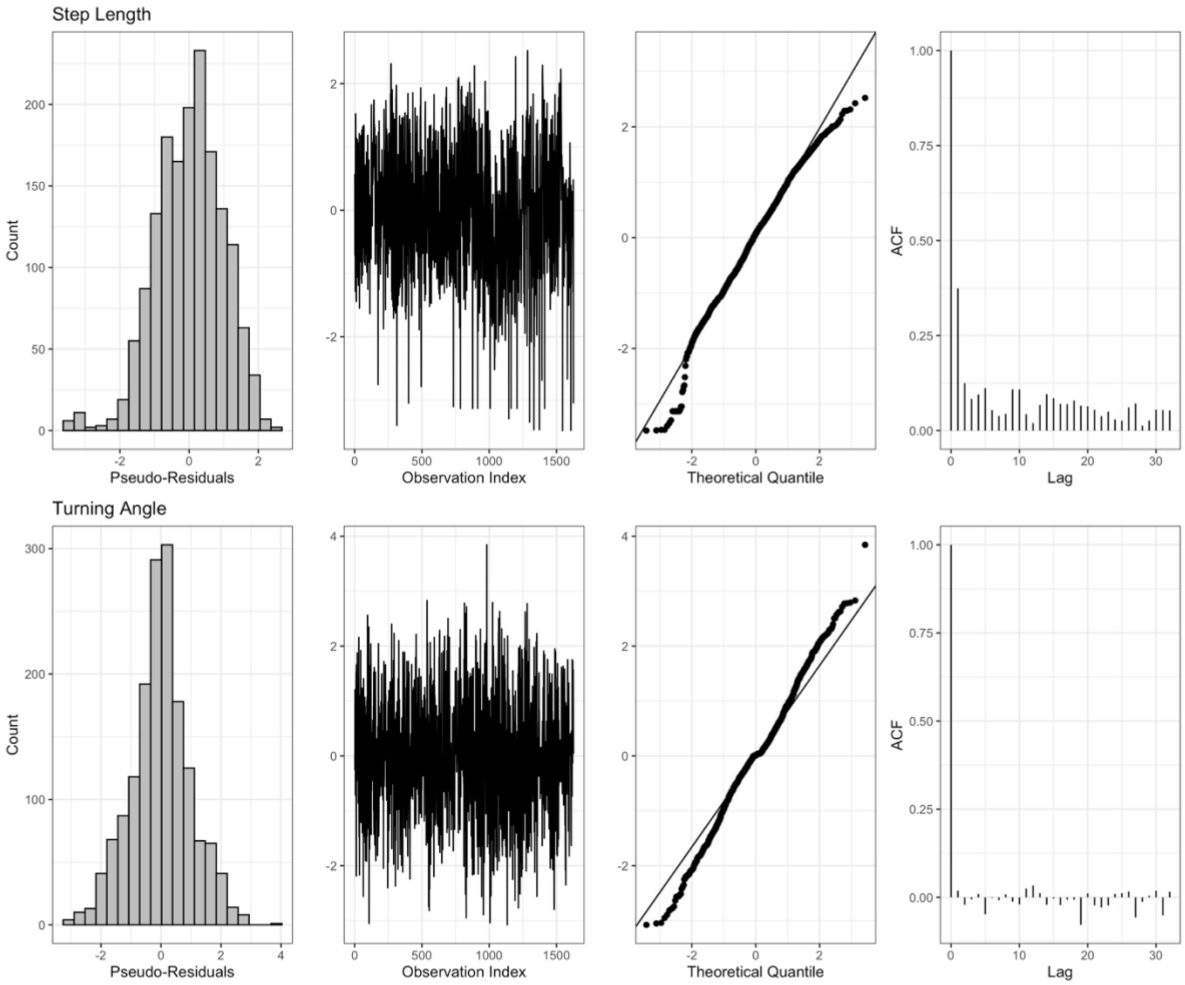


**Figure D1.** Pseudo-residuals from the semi-supervised HMM

Model evaluation was conducted by visually inspecting the pseudo-residuals. The Q-Q plot showed good agreement with normality in the central quantiles, especially for step length, but deviated from the reference line in the upper and lower tails. This, along with the ACF plot indicate some autocorrelation in step length is present but residual autocorrelation is expected to have only a minimal influence on behavioural state decoding.

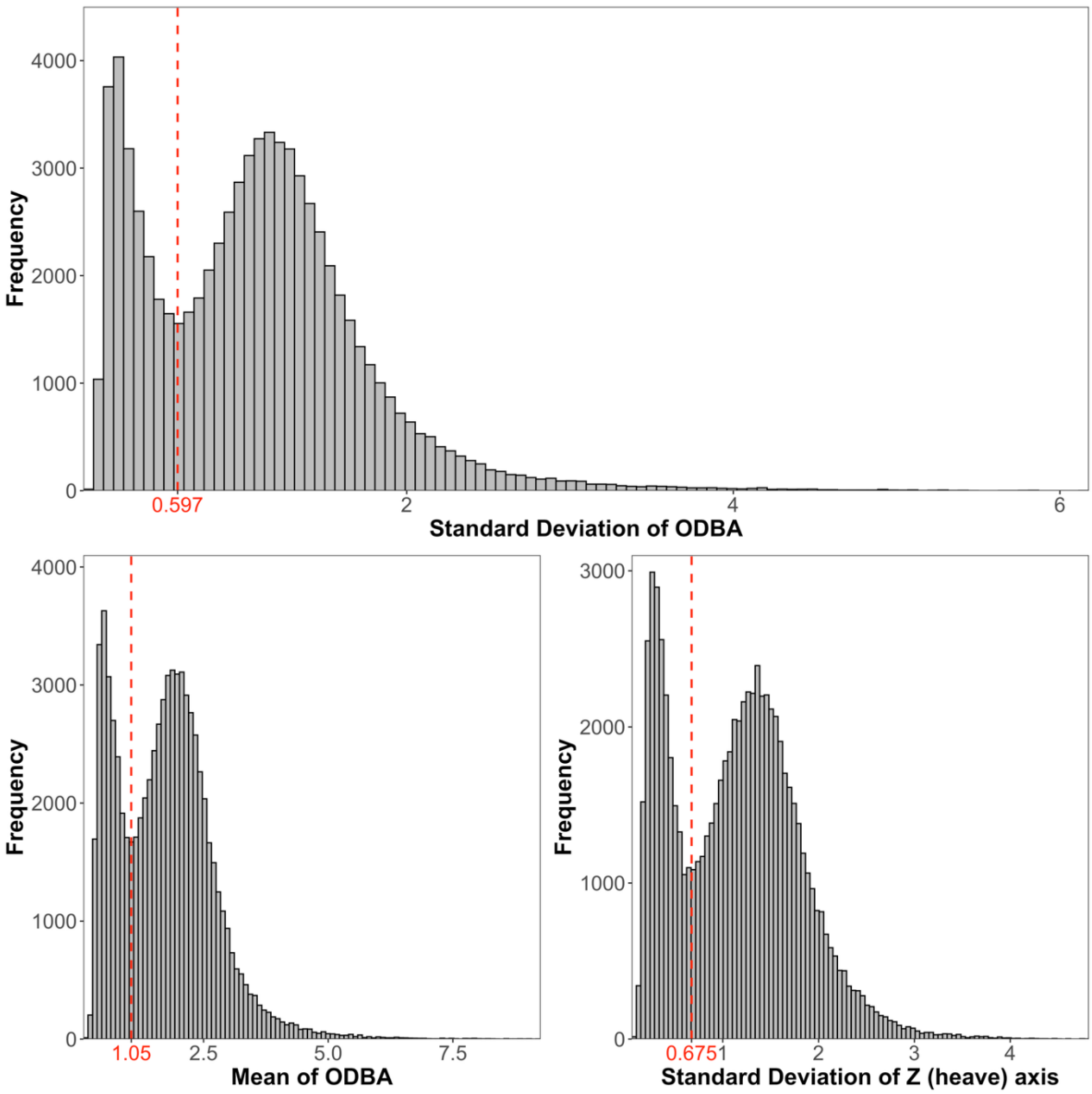


**Figure S1.** Histograms of the standard deviation of Overall Dynamic Body Acceleration (ODBA), mean of ODBA, and standard deviation of the Z (heave) axis for a European storm-petrel foraging trip. The red dashed line and x-axis label indicates the interpeak frequency minimum. The standard deviation of ODBA was used to separate low-activity (resting) and high-activity (flying) behaviours (resting is < 0.597 $SD_{ODBA}$). See Figure S2 for histograms for the other two accelerometer datasets.

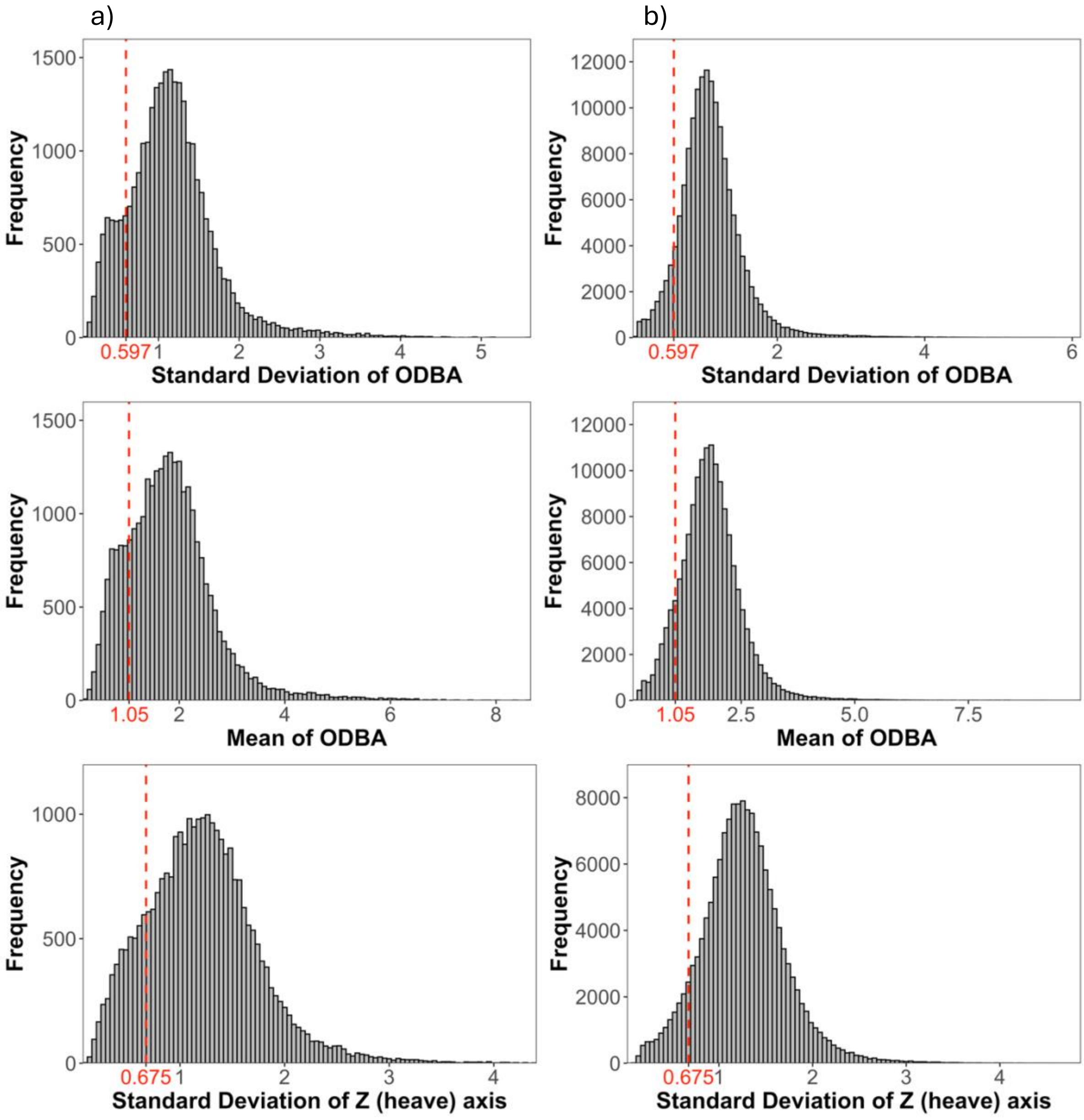


**Figure S2.** Histograms of the standard deviation of Overall Dynamic Body Acceleration (ODBA), mean of ODBA, and standard deviation of the Z (heave) axis for two European storm-petrel foraging trips with minimal resting behaviour. The red dashed line and x-axis label indicates the interpeak frequency minimum calculated from the trip featured in Figure S1.

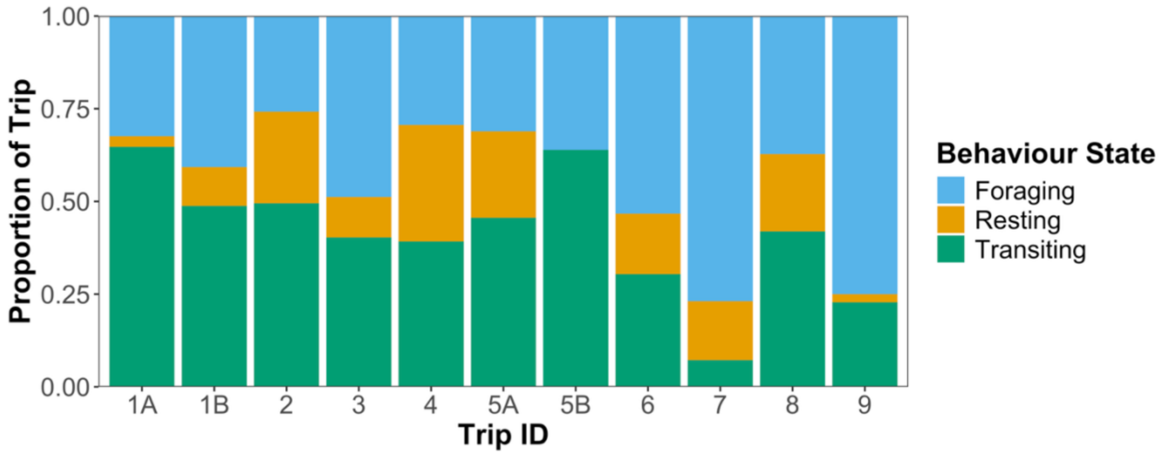


**Figure S3.** Proportion of GPS points classified as each behaviour state for the complete foraging trips (n = 11) according to the three-state HMM informed by accelerometer data. Two consecutive trips were recorded for individuals 1 and 5.

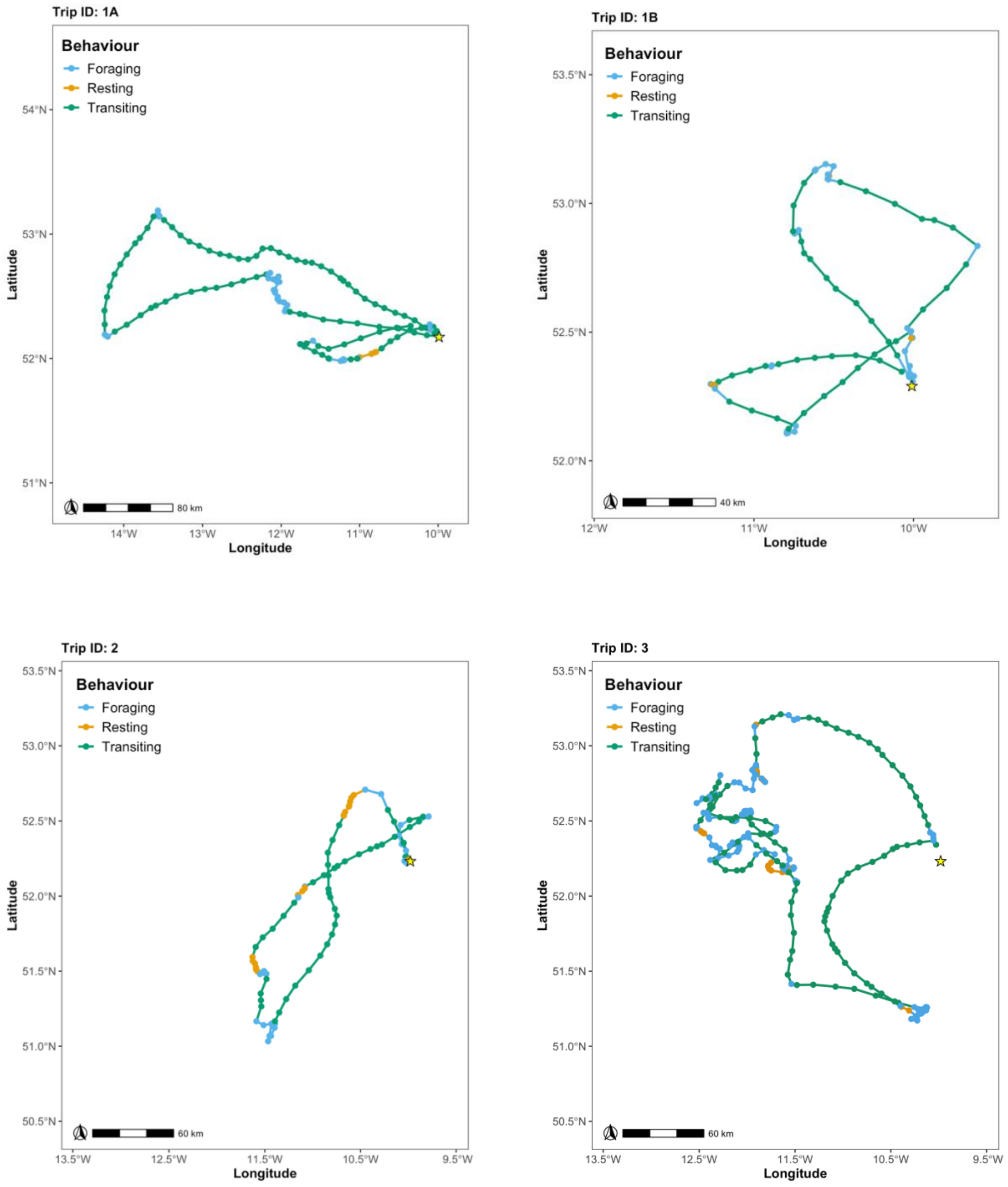


**Figure S4.** Foraging trips segmented into foraging (blue), resting (orange), and transiting (green) behaviours according to the accelerometer-informed three-state HMM. The breeding colony location is marked with the yellow star.

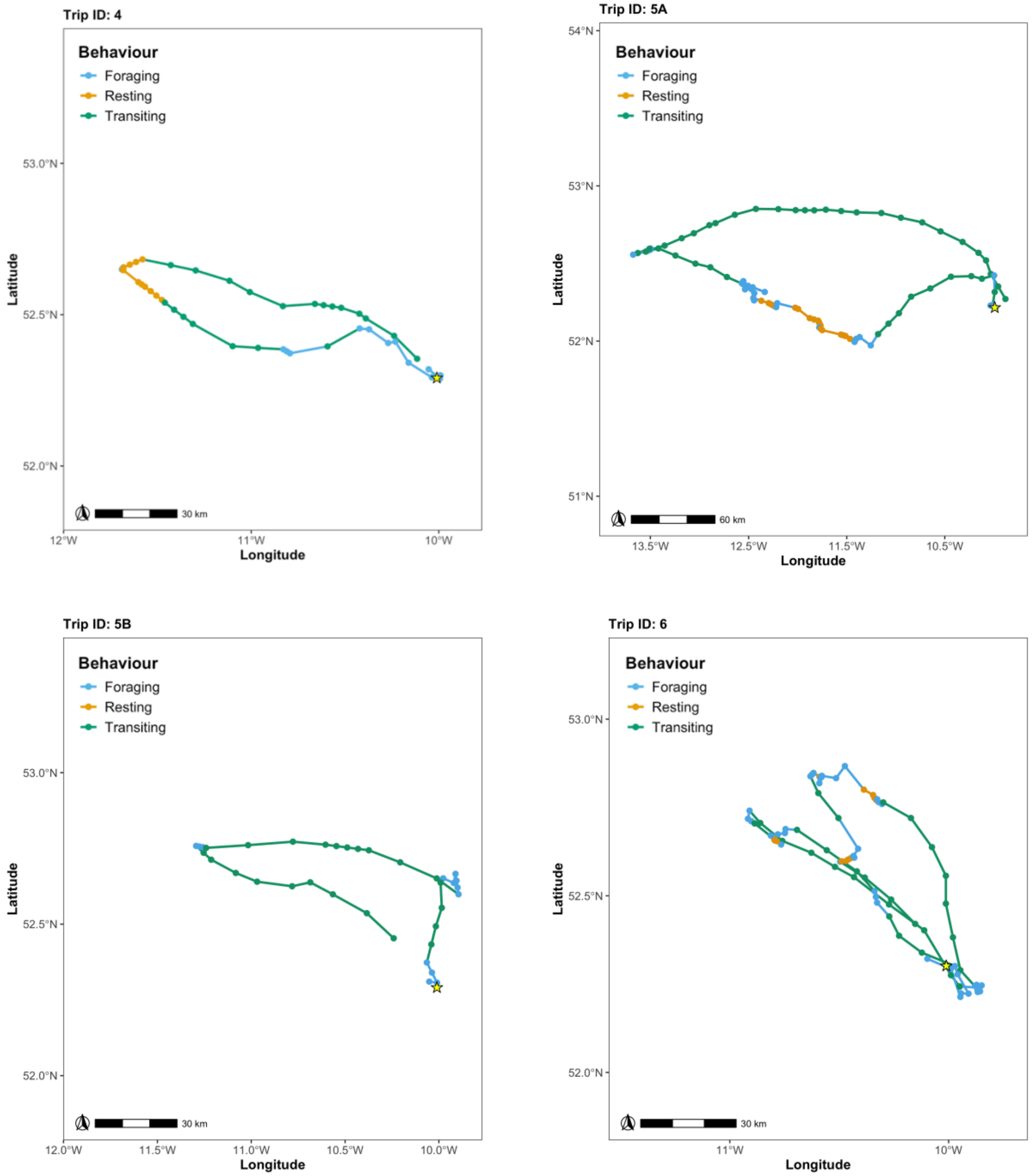


**Figure S4.** Continued

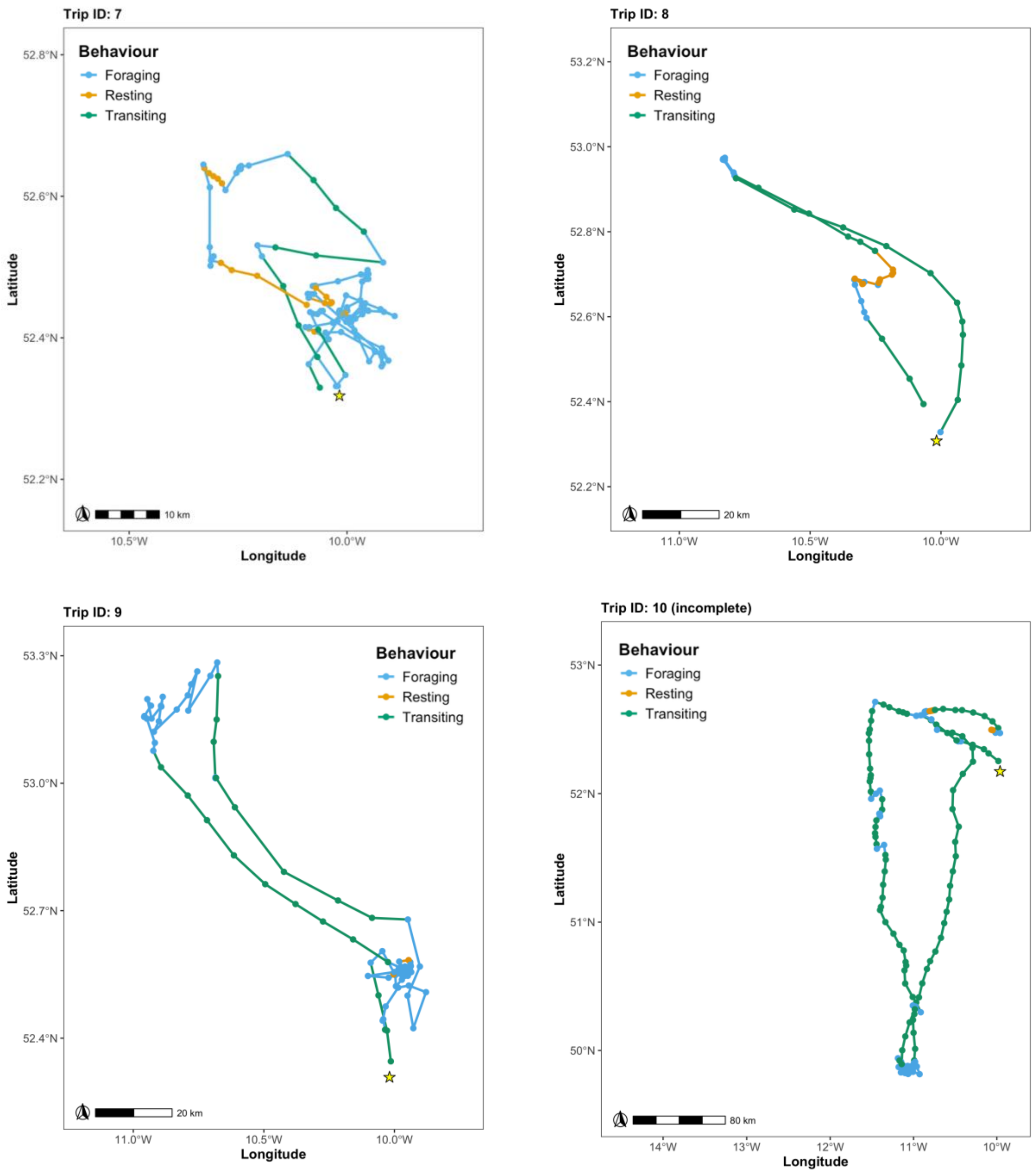


**Figure S4.** Continued

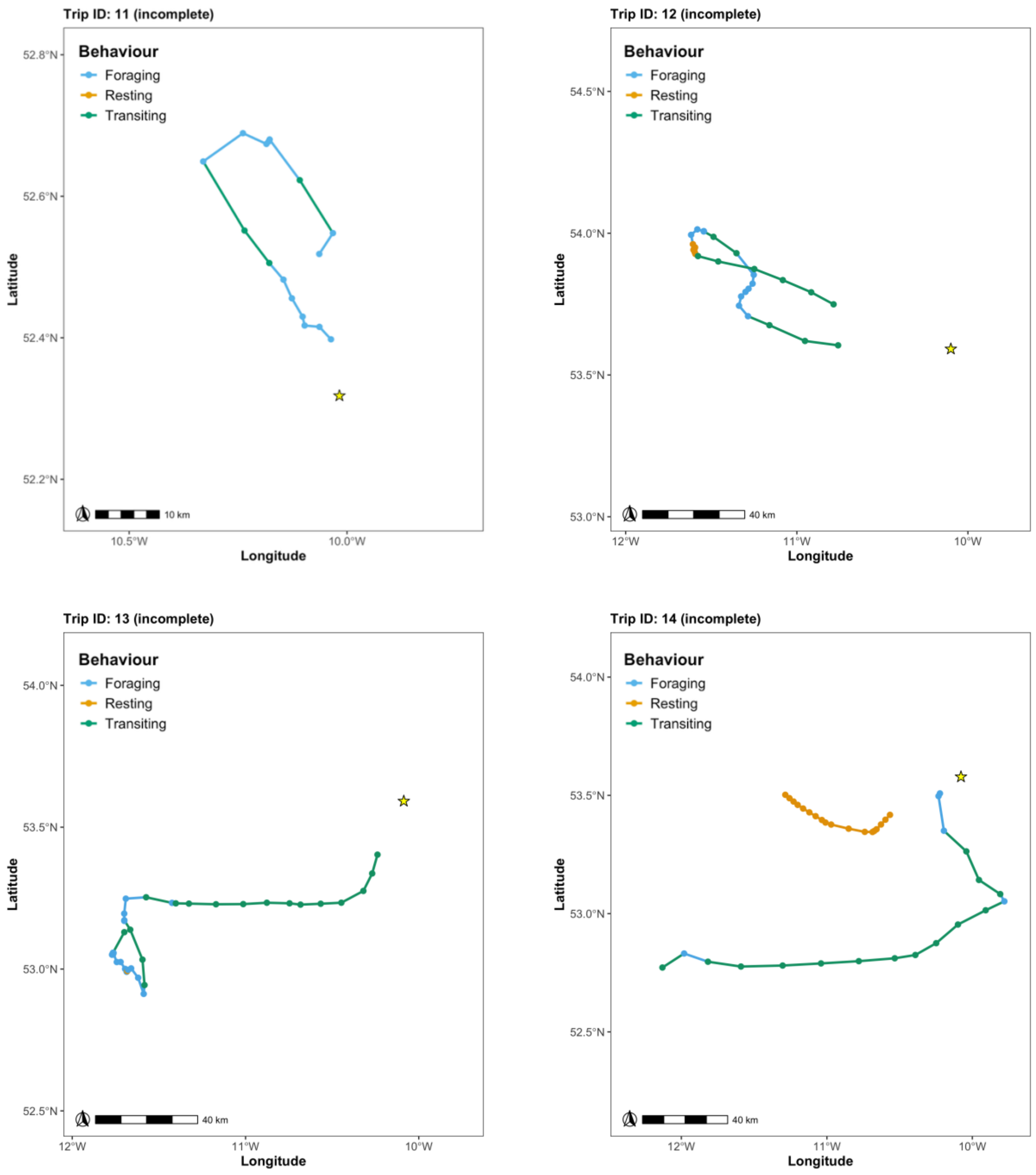


**Figure S4.** Continued

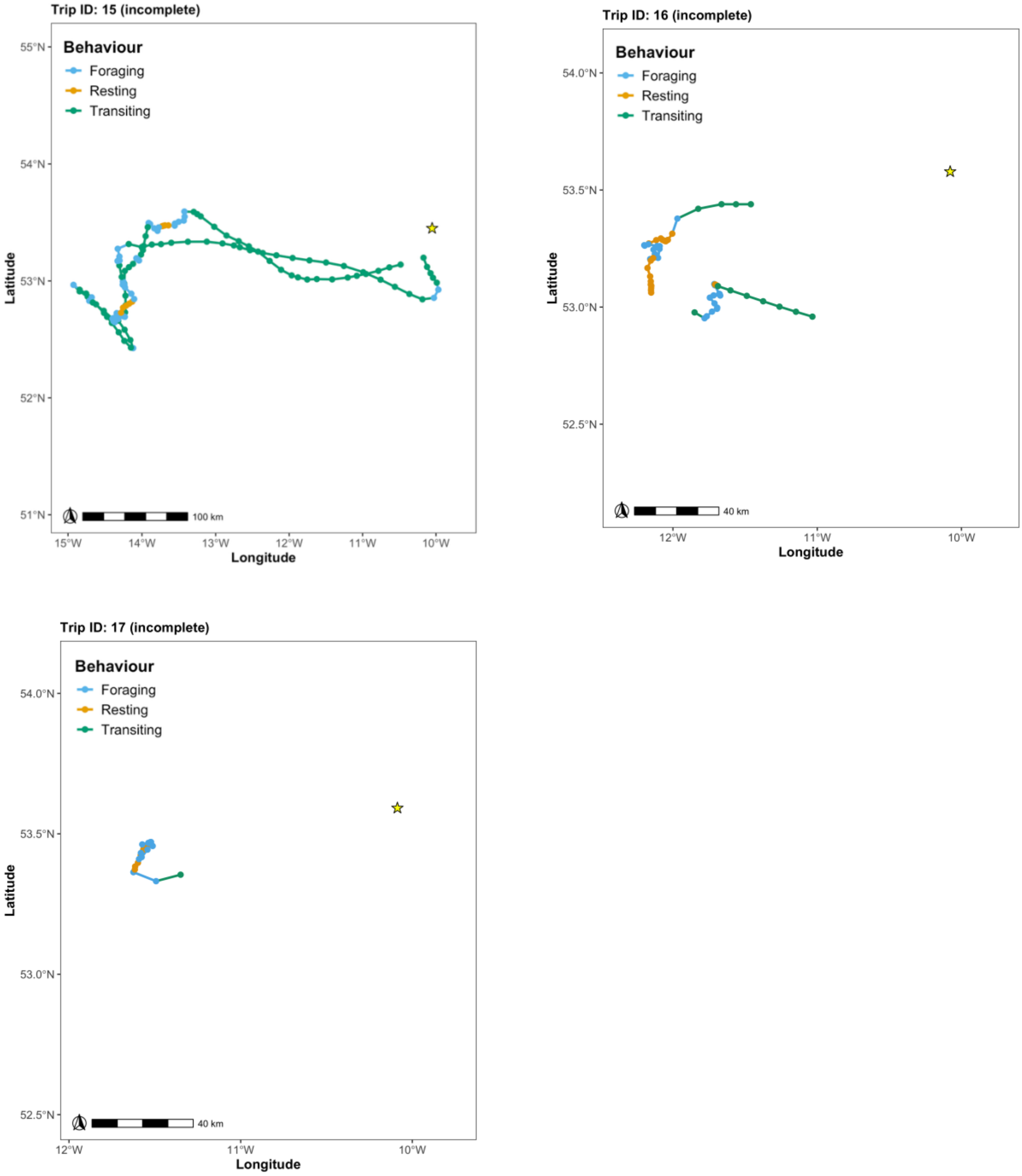


**Figure S4.** Continued

**Table S1.** Log-likelihoods, difference (Δ) in log-likelihood, Akaike Information Criterion (AIC), and Δ in AIC between Hidden Markov Models produced with 1-4 states. The largest Δ in log-likelihood was chosen as the method of selecting the most suitable number of states as the AIC often fails to select the correct number of states when applied to real data (Pohle et al., 2017).

| Behaviour states | Log-Likelihood | Δ Log-Likelihood | AIC | Δ AIC |
|---|---|---|---|---|
| 1 | 17989.5 | - | 35987.0 | - |
| 2 | 17087.7 | 901.8 | 34197.4 | 1789.6 |
| 3 | 16850.5 | 237.2 | 33741.0 | 456.4 |
| 4 | 16716.7 | 133.8 | 33495.3 | 245.7 |

**Table S2.** Confusion matrix of the number of GPS points classified as foraging, resting, and transiting by the accelerometer-informed HMM (rows) and the GPS-only HMM (columns). Diagonal cells (highlighted in green) show classification agreement.

| | Foraging | Resting | Transiting | Total |
|---|---|---|---|---|
| **Foraging** | 718 | 9 | 0 | 727 |
| **Resting** | 176 | 38 | 5 | 219 |
| **Transiting** | 34 | 13 | 612 | 659 |
| **Total** | 928 | 60 | 617 | 1605 |
